# Optical Cellular Micromotion: A New Paradigm to Measure Tumour Cells Invasion in 3D Tumour Environments

**DOI:** 10.1101/2021.08.26.457857

**Authors:** Zhaobin Guo, Chih-Tsung Yang, Chia-Chi Chien, Luke A. Selth, Pierre O. Bagnaninchi, Benjamin Thierry

## Abstract

Measuring tumour cell invasiveness through three-dimensional (3D) tissues, particularly at the single cell level, can provide important mechanistic understanding and assist in identifying therapeutic targets of tumour invasion. However, current experimental approaches, including standard *in vitro* invasion assays, have limited physiological relevance and offer insufficient insight about the vast heterogeneity in tumour cell migration through tissues. To address these issues, here we report on the concept of optical cellular micromotion, where digital holographic microscopy (DHM) is used to map the optical thickness fluctuations at sub-micron scale within single cells. These fluctuations are driven by the dynamic movement of subcellular structures including the cytoskeleton and inherently associated with the biological processes involved in cell invasion within tissues. We experimentally demonstrate that the optical cellular micromotion correlates with tumour cells motility and invasiveness both at the population and single cell levels. In addition, the optical cellular micromotion significantly reduced upon treatment with migrastatic drugs that inhibit tumour cell invasion. These results demonstrate that micromotion measurements can rapidly and non-invasively determine the invasive behaviour of single tumour cells within tissues, yielding a new and powerful tool to assess the efficacy of approaches targeting tumour cell invasiveness.

**Significance Statement:** Tumour cells invasion through tissues is a key hallmark of malignant tumour progression and its measurement is essential to unraveling biological processes and screening for new approaches targeting cell motility. To address the limitations of current approaches, we demonstrate that sub-micron scale mapping of the dynamic optical thickness fluctuations within single cells, referred to as optical cellular micromotion, correlates with their motility in ECM mimicking gel, both at the population and single cell levels. We anticipate that 3D optical micromotion measurement will provide a powerful new tool to address important biological questions and screen for new approaches targeting tumour cell invasiveness.

## Introduction

Metastasis is the main direct cause of tumour-related mortality. It is initiated by tumour invasion to nearby/distant organs as a result of tumour cells breaking the basement membrane barrier and migrating into adjacent tissues. Tumour cell invasion is therefore a key hallmark of malignant progression. Investigation of factors that drives tumour cell invasion has the potential to reveal new approaches to control early stage diseases^1, 2^ as well as to provide better understanding of the role of tumour cell phenotypes and tumour microenvironment on the metastatic process^3–6^. While a number of conventional cell invasion assays exists, they suffer from several limitations that negatively impact progress in this field. For example, commonly used wound healing or Boyden chamber assays are laborious and time consuming, typically requiring hours or days for data acquisition. Moreover, the readout of these assays are accumulative invasion effects between observation intervals, such as migration distance or coverage of wounds, which fail to inform on the transient invasiveness changes in response to external factors. More importantly, these assays do not recapitulate the 3D environment experienced by cancer cells in intricate contact with the extracellular matrix (ECM)^7–9^, which limits their relevance for the screening of potential activators or suppressors of tumour invasion. Finally, current invasion/migration assays are typically population-based, and thereby do not provide insights about the role of tumour cells heterogeneity in these processes.

The invasion of tumour cells within tissues is inherently associated with cellular structural changes, including the formation of lamellipodia, filopodia, and integrin-mediated adhesion^10, 11^, in which the cytoskeleton (e.g. actin, myosin, etc.) plays a pivotal role. This led us to hypothesize that, independently of the upstream effectors of invasion, the invasiveness of a specific tumour cell within its 3D microenvironment can be directly correlated to the cellular micromotion. Cellular micromotion refers to the biologically-driven continuous nanoscale movement of the cell membrane and intracellular structures, including the cytoskeleton.^12–15^ Micromotion can be experimentally measured using electrical impedance changes associated with nanoscale to submicron scale fluctuations of the membrane of living cells on microelectrodes, as initially described by Giaever and Keese^12^. In support of this early pioneering work, several studies have since shown that cellular micromotion as measured by electrical impedance can not only identify cancerous cells but also provide insight into their metastatic potential^13, 15, 16^. In addition, phenotypic changes linked to invasion such as epithelial mesenchymal transition (EMT), are also associated with measurable changes in cellular micromotion^17, 18^.

While the monitoring of cellular micromotion therefore provides a powerful approach to measuring, within a few minutes, the invasiveness of tumour cells, the current *modus operandi* based on impendence measurements suffers from several conceptual and practical limitations. The monitoring of micromotion by impedance requires the formation of a cell monolayer onto the solid substrate serving as electrode. While this makes for an easy experimental set-up, it presents limited physiological relevance to the invasiveness of cells within the 3D environment of real tissues. In addition, it does not allow for single-cell measurements and consequently provides no insights into the inherent heterogeneity of the invasion process.

To validate our hypothesis that cell invasiveness has a significant correlation with cell micromotion, and to overcome the shortcomings of impedance-based measurements of the micromotion, we endeavoured to measure the micromotion of individual cells within a 3D environment by monitoring and processing the spatiotemporal fluctuations of the optical thickness. To this end, we utilized digital holographic microscopy (DHM) to record the instantaneous optical thickness fluctuations of single cells embedded within fibrin gels and developed a dedicated algorithm to obtain a 3D micromotion index from these measurements. To demonstrate the link between the 3D micromotion index and invasiveness, the breast cancer cell line MDA-MB-231 was treated with epidermal growth factor (EGF) to induce a more invasive phenotype. The micromotion indexes of the treated cells suspended in a fibrin gel were significantly increased in comparison to the untreated ones. Further, we demonstrate that the 3D micromotion index is correlated at the single cell level to their motility within the fibrin gel measured as the mean migration speed over 12h. To further validate our hypothesis, we showed that cells in more invasive mesenchymal phenotypes also displayed significantly increased 3D micromotion indexes. A direct correlation was also measured at the single cell level between vimentin expression and micromotion for A549 cells. In addition, the micromotion of miR-194 overexpressing PC-3 cells was measured and compared to that of normal PC-3 cells. We have previously shown that miR-194 is a driver of prostate cancer invasiveness and stable overexpression enhanced metastasis of intravenous and intraprostatic tumor xenografts^19^. MiR-194 PC-3 cells were also found to have significantly higher extravasation rates in a microfluidic vasculogenesis model and yielded significantly higher micromotion than that of non-transfected cells. Finally, PC-3 cells treated with cellular motility targeting (migrastatic) compounds that have been shown to inhibit prostate cancer cells invasion and metastasis^20, 21^ yielded significantly reduced micromotion, and the dynamic of micromotion alteration is in good agreement with the effectiveness of compounds in invasion inhibition. Altogether, this data validates the concept that a 3D micromotion index obtained from dynamic phase measurements provides rapid and simple measurement of tumour cells invasiveness in the 3D environment at the single cell level, and shows the feasibility of using optical 3D cellular micromotion to investigate the effectiveness of novel migrastatic approaches. The application of this new paradigm could accelerate the screening for novel therapies as well as assist in addressing important biological questions about the metastatic process.

## Results

### Probing and quantifying optical cellular micromotion in 3D environments

We first endeavoured to demonstrate the feasibility of measuring the dynamic optical thickness (OT) fluctuations of cells within a 3D environment mimicking that of tumour tissues. We selected fibrin gel as the model 3D extracellular matrix due to its inherent high bioactivity and suitability for studies related to cellular adhesion and degradation processes. Fibrin gel promotes typical invasive behaviours, for instance microenvironment remodelling and migration and is commonly used in advanced tumour models, both *in vivo* and *in vitro*^12, 23^. MDA-MB-231 breast cancer cells were embedded in 5 mg/mL of fibrin gel and loaded in a custom-made microfabricated device, in which both cells and gels are sandwiched between two coverslips. This configuration provides a homogeneous optical background required for high precision phase imaging with DHM. Holographic movies of single cells in the fibrin gels were captured and converted to quantitative phase images (Fig. 1A and SI movie 1), from which the phase shifts were extracted and segmented from background (Fig. 1B-1D). DHM requires only low intensity illumination, and it is therefore possible to perform imaging continuously at high capture frequencies and/or repeatedly. In our experimental set-up, high frequency acquisition enabled the capture of the fluctuations of the OT for whole cells (Movie S1).

**Figure 1.**
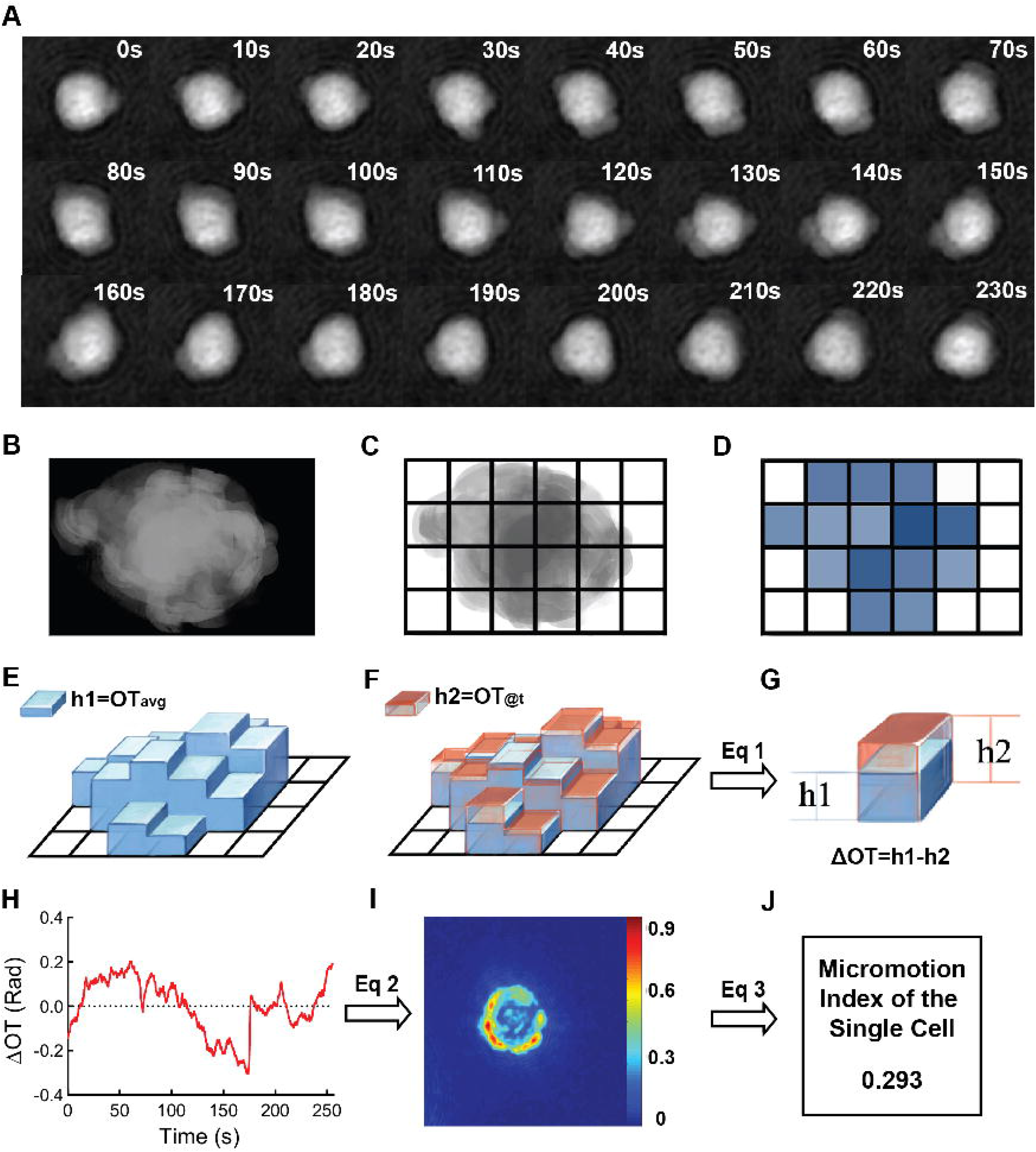
Probing and quantifying optical cellular micromotion in 3D environments. **(A)** Time sequence image series of single cell in fibrin gel imaged with DHM. **(B-J)** Schematic illustration of the work flow for single cell micromotion measurement. **(B)** DHM imaging of a single cell **(C)** Segmentation of cell from background. **(D)** Pixel based histogram plotting of mean optical thickness (OT) over entire image period. **(E)** Pixel based histogram of mean OT over entire image period. **(F)** OT at specific time point T_a_. **(G)** Calculation of Δ OT (OT@T_a_-Mean OT) using Eq1. **(H)** Plot of ΔOT fluctuation over time. **(I)** Calculated mean square of fluctuation (micromotion index) using Eq 2 and mapping over entire cell. **(J)** Averaging micromotion index of all pixels in the cell.

Micromotion indexes have been previously obtained by calculating the mean square of the stochastic signal fluctuations associated with the presence of live cells on the solid substrates ^24–26^. In the proposed optical micromotion paradigm, one can either consider the OT fluctuations for an entire cell (e.g. averaging the fluctuation of all the pixels comprised within the cell) or the fluctuation of every pixel within a single cell. To illustrate the difference between the two approaches, we randomly selected 3 pixels inside a cell and plotted the fluctuations of the associated OT over time. We also plotted the mean OT calculated for the same whole cell (Fig. S1). From these measurements, it is clear that the process of averaging results in significant dampening of the OT fluctuations. This is not surprising as over the short period of time of these measurements, the overall OT of a cell is relatively constant. On the other hand, biological processes result in significant redistribution of the dry mass of ‘optically thick’ intracellular structures, which can be detected when monitoring single pixels. Dry mass refers to the mass of all the substances within the cells (e.g. protein, nucleic acid, etc.) other than water. We therefore quantified the dynamic fluctuation of the OT for every pixel inside each cell (Eq. 2), and then averaged these values over the whole cell (Eq. 3) to obtain the single cell MI. The entire measurement process is illustrated in Fig. 1E-J.

### Measured 3D optical cellular micromotion are mainly associated with live cell behaviours

To confirm that the calculated MI is associated with biological processes, we compared values obtained for live cells with those obtained for fixed cells, both in the 3D and 2D (cells plated on a coverslip) environment. As shown in Fig. 2A and B, significantly lower fluctuations were observed for fixed cells both in 2D and 3D. The calculated MI indexes of 30 individual cells are presented in Fig. 2C. The MIs of live MDA-MB-231 cells ranged from 0.07 to 0.27 (average of 0.165) in the 3D environment while the MIs of fixed cells ranged from 0.02 to 0.04 (average of 0.032). Similarly, the averaged MI for live cells in 2D was 0.07 while the average MI for fixed cells was 0.018. This data confirm that the measured dynamic fluctuations are mainly associated with live cell rather than to instrumental noise or physical perturbations. The MI was also calculated at 6h or 24h after being seeded into the 3D fibrin gel, and no significant differences were found neither for the MDA-MB-231 nor MCF-7 cells (Table. S1).

**Figure 2.**
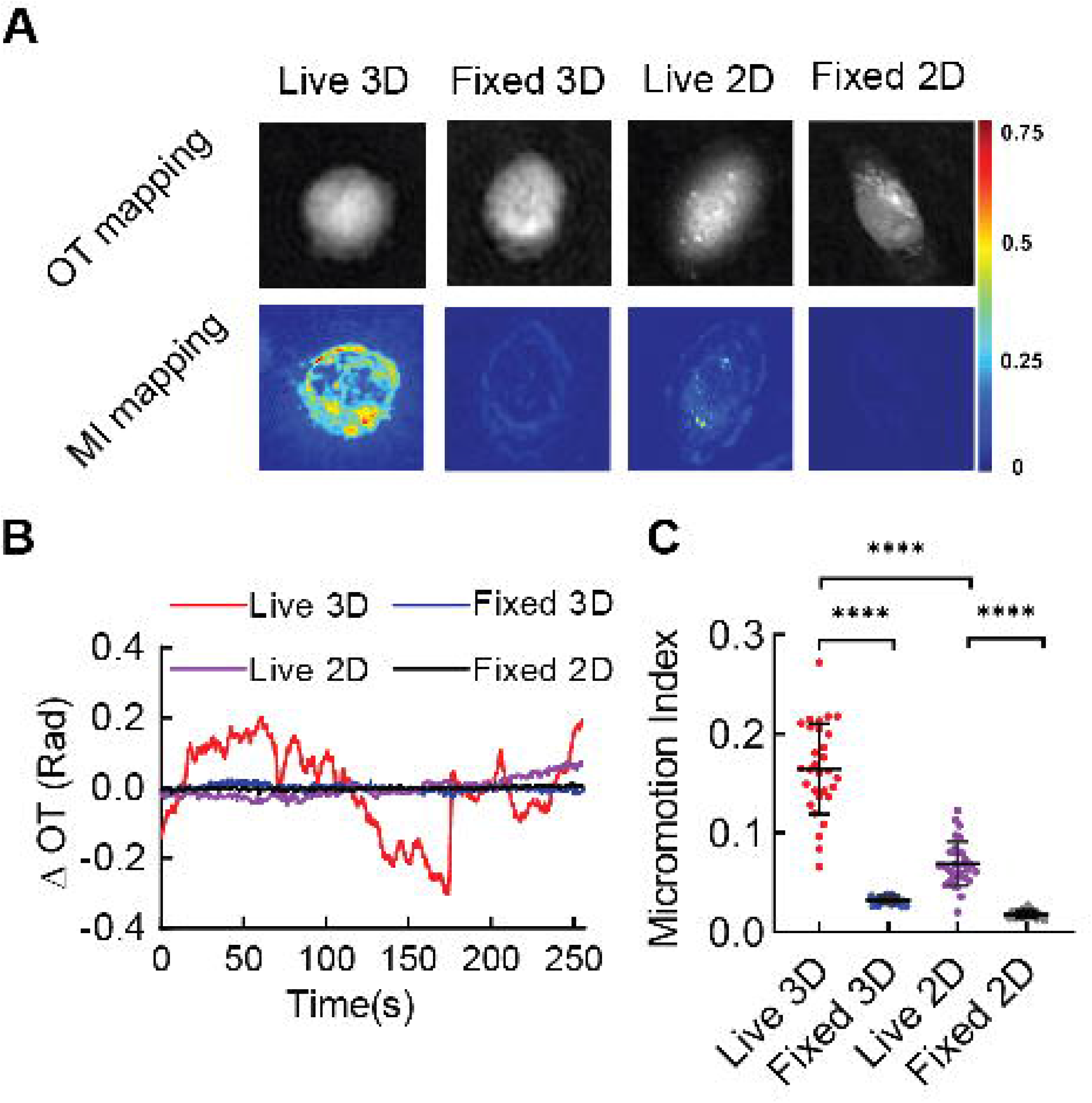
Measured 3D cellular micromotions are mainly associated with live cell behaviors and significantly higher than in 2D. **(A)** Fluctuation curves of live and fixed MDA-MB-231 cells in 2D and 3D (at 24h). **(B)** OT and MI mapping of live and fixed cells in 3D and 2D. **(C)** MI measurements for live and fixed cells in 3D and 2D (one point represents the MI of a single cell). Error bars are SD (****, P< 0.0001 unpaired two-sided t-test)

### Measured 3D optical cellular micromotion are significantly higher than that in 2D

Our initial experiments with the MDA-MB-231 model also allowed us to directly compare the cellular micromotion in 2D and 3D environments (Fig. 2A and B, Movie S1 and S2). The mean MI of the MDA-MB-231 cells in 3D fibrin gel was significantly higher than that of cells plated on a cover-slip (2D) (0.165 ± 0.045 vs 0.069 ± 0.021, p< 0.001). Meanwhile, the minor MIs difference between fixed cells in 3D and 2D (0.032 ± 0.004 vs 0.018 ± 0.004) indicates that the significant differences observed for the live cells between the 2D and 3D environments indeed originates from biological phenomena instead of experimental or technical discrepancies. While the reason for the difference of MIs between in 2D and 3D is not clear, it likely originates in the increased rigidity of cells adhered to aberrantly hard substrates which limits cell membrane and intracellular motility. On the other hand, it is also likely that cells inherently display higher mechanobiological activities or different modality of invasion when in a 3D environment mimicking that of real tissues^27, 28^.

### Micromotion and invasiveness in fibrin gel are correlated at the single cell level

We next investigated whether micromotion measurements could be correlated to the invasiveness of cells within a 3D environment. To modulate the invasive capability of MDA-MB-231 cells, we treated cells with the epidermal growth factor (EGF), which increases invasiveness *in vitro* and *in vivo*^29, 30^. The MDA-MB-231 cells were dispersed within a 3D fibrin gel supplemented or not with 100 ng/mL EGF. The micromotion indexes of the MDA-MB-231 cells was then measured from the DHM as described above at 6h after seeding (Fig. 3A). Following DHM imaging, these cells were individually tracked to establish their migration paths in the following 12h, taking an image every 10 min. Herein we use the motility to reflect the invasiveness of cancer cells, since these two parameters are closely related and motility can be easily tracked in this setting. Consistent with what has been reported previously, treatment with EGF significantly increased the motility of MDA-MB-231 cells in fibrin gel (Fig. 3B). This is quantitively demonstrated by differences in the ‘mean migration speed’, which refers to the mean migration distance between 2 consecutive frames (Fig. 3C) (mean speeds of 1.74 ± 0.91 μm/10 min for +EGF vs 1.17 ± 0.47 μm/10 min for control, p< 0.01**). The mean MIs calculated from 5 min DHM measurements for these two cellular populations were also significantly different (Fig. 3A, 0.248 ± 0.140 for the +EGF group vs 0.170 ± 0.056 for the control group, p< 0.01**).

**Figure 3.**
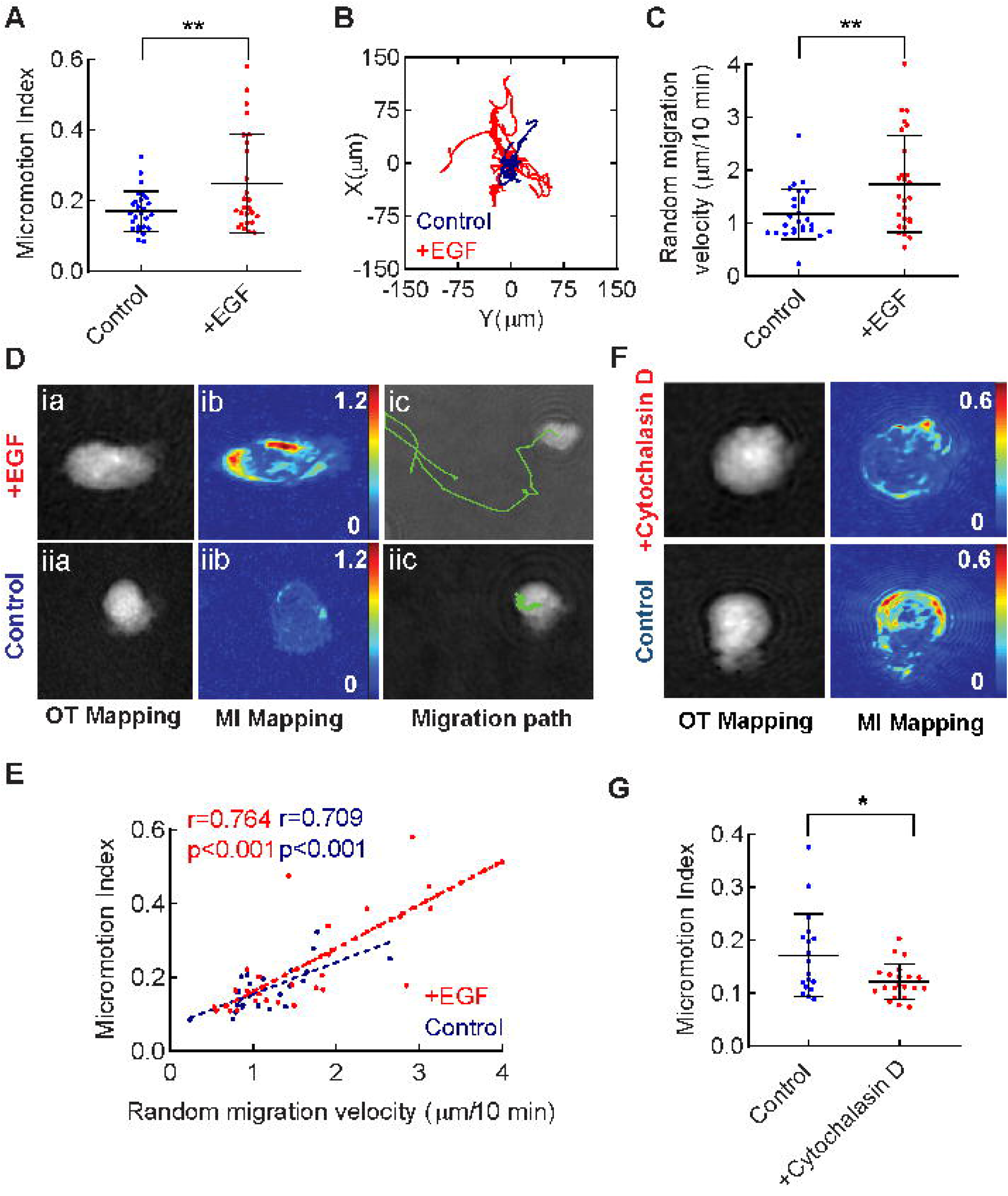
Micromotion and motility in fibrin gel are correlated at the single cell level. **(A)** MIs measured for MDA-MB-231 cells treated or not with EGF. **(B)** Typical migration paths recorded over 12h for two single cells within fibrin gel supplemented or not with EGF. **(C)** Mean migration speeds in fibrin gel calculated for single cells treated or not with EGF (2D projection). **(D)** OT mapping, MI mapping and migration tracking of a typical cell in the +EGF (**Dia-Dic**) and control (**Diia-Diic**) groups. **(E)** Scatter plot of the MI vs mean migration speed for single cells in +EGF (red) and control (blue) groups. **(F)** OT mapping and MI mapping of a typical cell in the +Cytochalasin D and control groups. **(G)** MIs measured of cells treated or not with Cytochalasin D. Error bars are SDs (*P<0.05, ** P< 0.01, unpaired two-sided t-test)

From our results, it was also evident that there is significant heterogeneity in motility within the fibrin gel at single cell level, which is independent of treatment with EGF. This is not surprising as it has long been recognized that individual tumour cells respond differently to environmental stimuli due to inherent cellular heterogeneity at the genetic, epigenetic and phenotypic levels^31, 32^. This heterogeneity also reflects the observation that only a minority of highly invasive cells play central roles in tumour invasion^31^. Therefore, having demonstrated the correlation between the 3D cellular motility and MI at the population level, we then investigated whether single cell insights could be obtained. Fig. 3D display the OT mapping, micromotion mapping and migration path of a typical high MI cell in the EGF+ group while 3D-ii display those of a typical cell with a low MI (the micromotion and migration movies of corresponding cells are shown in Movie S3-S6). We plotted the MIs of single cells in both the EGF treated (Fig. 3E-i) and untreated (Fig. 3E-ii) groups vs their mean migration speeds determined over 12 h as described above. Pearson correlation (r) analysis demonstrated a strong correlation between the single cell micromotion and motility both for the EGF treated (r=0.764) and EGF untreated (r=0.709) groups.

Subsequently, we investigated the mechanisms underlying the correlation between cell motility and micromotion. More specifically, we studied the role of the actin cytoskeleton in regulating cellular micromotion, which is known as a pivotal regulator of tumour cell invasion. MDA-MB-231 cells were treated with a F-actin polymerization inhibitor cytochalasin D, after which micromotion indexes were measured. OT images and micromotion mappings of representative cells treated or not with cytochalasin D (+cytochalasin D and control group) are shown in Fig. 3F. A clear reduction of the MI was observed from the MI heat map of +cytochalasin D treated cell. Further statistical comparison of MI between both groups (18 from control and 21 from treated group) confirmed that the MI of cells was significantly reduced after cytochalasin D treatment (0.172 ± 0.076 vs 0.122 ± 0.033, P<0.05, Fig. 3G). Measurements in an invasive prostate cancer cell lines (PC-3) replicated the findings from the MDA-MB-231 model, although the reduction of MIs in prostate cancer cells was less pronounced (0.155 ± 0.059 vs 0.131 ± 0.033, P<0.05, Fig. S3). These sets of data support our hypothesis that the actin cytoskeleton plays a significant role in regulating tumour cell micromotion as defined in this work.

### Acquisition of EMT phenotype results in increased cellular micromotion

The acquisition of epithelial mesenchymal transition (EMT) phenotypes by tumour cells is associated with disruption of cell–cell adhesion and polarity, modulation in tumour cell–matrix adhesion, and remodelling of the cytoskeleton as well as with increased motility and invasiveness^33^. We therefore endeavoured next to determine whether the acquisition of an EMT phenotype is associated with increased optical cellular micromotion at the single cell level as previously observed in 2D^17, 18^. To address this question, we first treated the MCF-7 cell line with an EMT induction medium for 5 days. As shown in Fig. S2, compared to the non-treated group the EMT induced MCF-7 cells displayed significantly higher MIs (0.219 ± 0.097 vs 0.128 ± 0.043, P<0.001***), which confirmed that the transition of MCF-7 cells to a more invasive phenotype also led to increased micromotion. Next, to fully characterize how closely epithelial to mesenchymal phenotype transition and micromotion are associated, the A549 VIM RFP cell line was employed. This is a variant of the A549 cell line where vimentin is constitutively tagged with the red fluorescent protein (RFP), thereby enabling direct monitoring with confocal microscopy of the expression levels of this EMT marker. In this setting, we can use the fluorescence intensity of individual A549 cells to determine relatively the degree of phenotype transition. Consistent with the MCF-7 data, EMT induced A549 VIM RFP cells displayed significant higher MI values over the control group (Fig. 4A, 0.192 ± 0.064 for the EMT group vs 0.123 ± 0.033 for the control group, p< 0.001***). As expected and shown in Fig. 4B and C, the fluorescence intensities of EMT induced A549 VIM RFP cells were significantly higher than that of the untreated control group.

**Figure 4.**
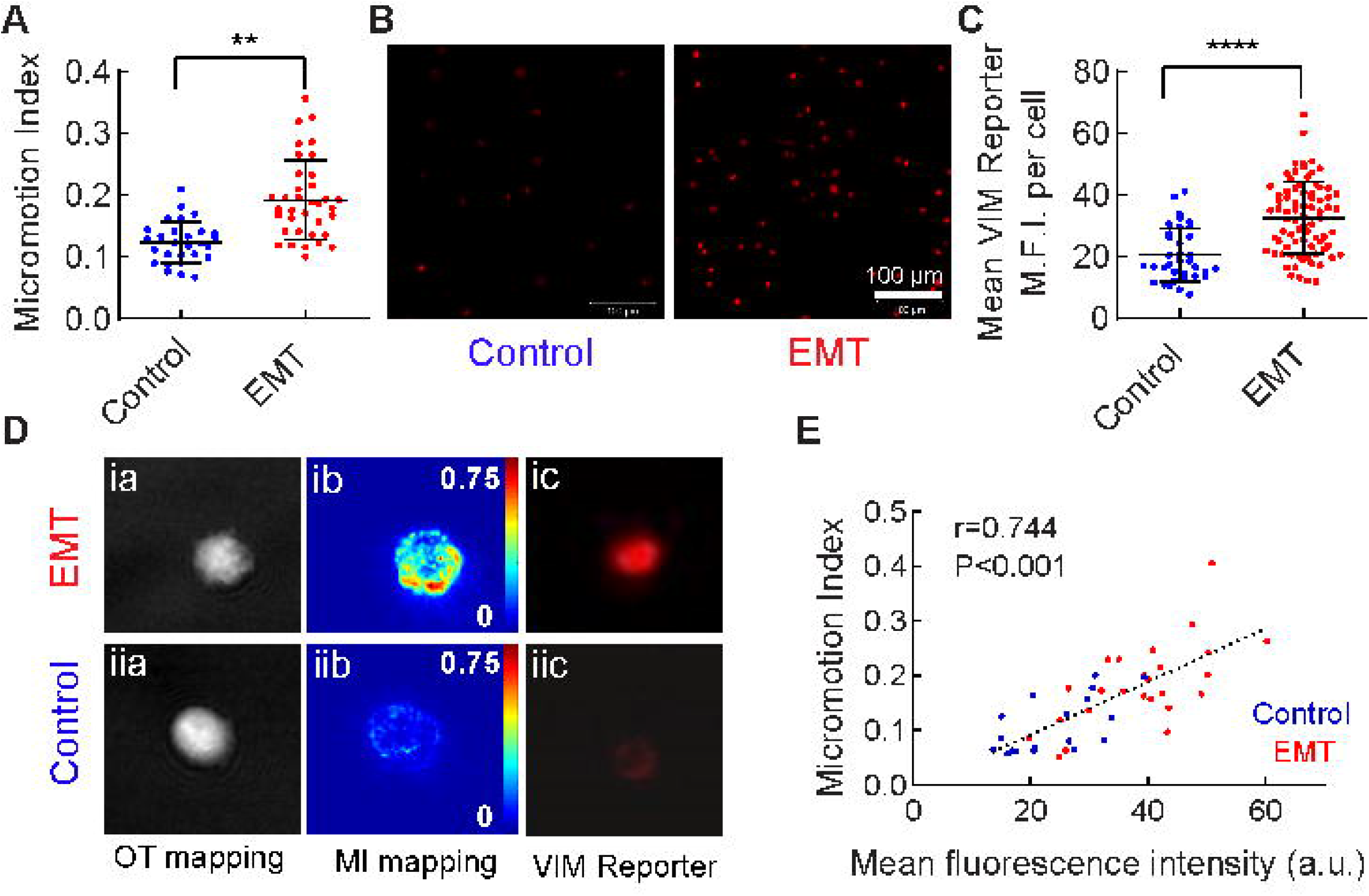
Acquisition of EMT phenotype results in increase in cellular MI. **(A)** MIs for EMT and control groups of A549 VIM RFP cells. **(B)** Confocal imaging of A549 VIM-RFP cells within fibrin gel with and without EMT induction. Scale bar: 100 μm. **(C)** MFIs for EMT and control A549 VIM RFP cells **(D)** Representative OT mapping, MI mapping and RFP imaging for A549 VIM-RFP cells with **(D-i)** or without **(D-ii)** EMT treatment. **(E)** Scatter plot of the MIs vs mean RFP fluorescence intensity for single cells in the EMT (red) and control (blue) groups. Error bars are SD (***, P< 0.001 unpaired two-sided t-test).

Next, we investigated whether the correlation exists at the single cell level. We plotted the micromotion (measured with DHM) over their RFP fluorescence intensity (measured with confocal microscopy) for every single cell, and calculated the correlation coefficient between these two quantities. Typical data for the OT mapping, micromotion mapping and fluorescence images of single cells from the EMT induction and control groups are presented in Fig. 4D-ia to ic and 4D-iia to iic, respectively. The mean fluorescence intensity of each cell was plotted vs the MI in Fig. 4E. From the data, a good correlation was obtained with a Pearson correlation coefficients (r) between these 2 factors of 0.744.

### Highly metastatic cells that display higher micromotion can be reduced by migrastatic compounds

Next, we investigated whether tumour cells specifically modified to display higher metastatic potential also exhibited higher micromotion within a 3D environment. We have previously shown that miR-194 promotes prostate cancer invasion and metastasis^19^. After being injected into the tail veins of mice, PC-3 cells overexpressing miR-194 have a much greater capacity to colonise organs and growth as assessed by whole-animal bio-luminescent imaging (Fig. 5A and B). To confirm this finding in an advanced *in vitro* setting, a microfluidic perfusable model of the human microvasculature was used. In this model, live cancer cells can be perfused within the ‘on-chip microvasculature’ to investigate the trans-endothelial migration/extravasation. This model has been previously successfully used to investigate the invasiveness of different cell lines^34–36^. Fig. 5C shows typical images for the miR-194 overexpressing and control groups 24h after being perfused in the microvasculature. The extravasation ratios (number of extravasated cells/total number of cells) were calculated from randomly selected images. In agreement with the *in vivo* data, miR-194 overexpressing PC3 cells displayed significant higher extravasation in comparison to the control group at 24h. A trend for higher extravasation was also measured at 9h for the miR-194 over-expressing group as shown in Fig. 5D. Subsequently, we measured the micromotion of miR-194 overexpressing PC-3 cells dispersed within the fibrin gel. MiR-194 over-expressing PC-3 cells displayed significantly higher micromotion indexes compared to that of the control cells (Fig. 5E and F) (0.199 ± 0.061 vs 0.153 ± 0.041, p< 0.001***), providing further support that this measurement can be used as a proxy for invasion and metastasis.

**Figure 5.**
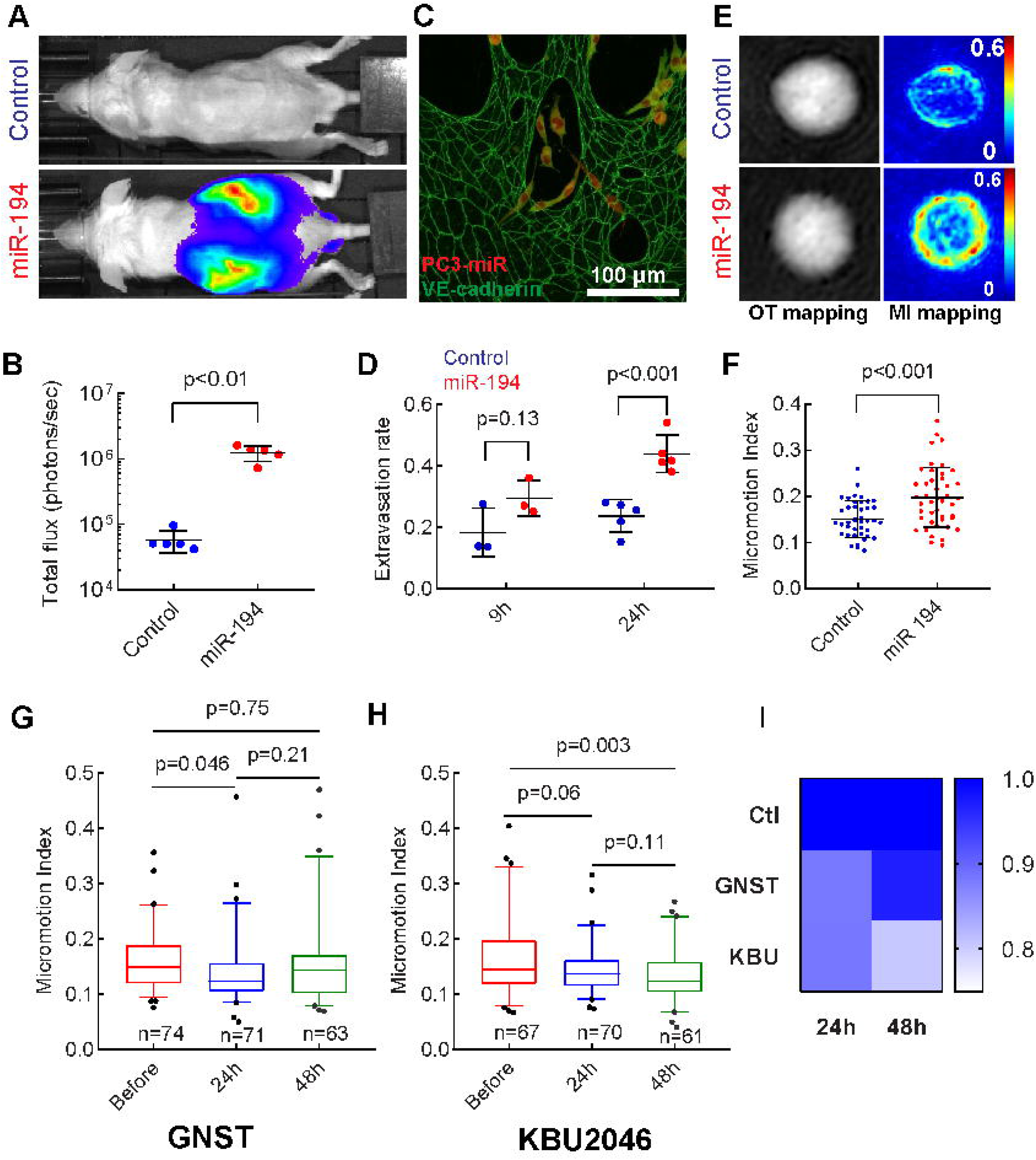
Highly metastatic cells that display higher optical cellular micromotion can be reduced by migrastatic drugs. **(A)** Images showing whole-animal bioluminescent imaging of representative mice in each group at 6 weeks after mice were injected with control cells or miR-194-overexpressing cells. **(B)** Luciferase intensity as assessed by whole-animal bioluminescent imaging (n=5 in each group, one data point represents a single mouse). **(C)** Confocal imaging of PC-3 cells extravasation in a microfluidic microvasculature model. Vasculature is stained for VE-cadherin (green) and PC-3 are labelled with CellTracker™ CMTPX dye (red). **(D)** PC-3 cells extravasation ratios in the microfluidic microvasculature model for non-transfected control (black) and miR-194 overexpressing cells (red) at 9h and 24 after perfusion. One point represents a separate microfluidic device. **(E)** Typical images of PC-3 cells OT and MI mapping. **(F)** MIs for miR-194 overexpressing PC-3 cells and control group. **(G-H)** MIs of PC-3 before, 24h and 48h after being treated with 10 μM of genistein or KBU 2046. Error bars are SD (unpaired two-sided t-test). **(I)** Fold changes of mean MIs of PC-3 cells at different time points after being treated with genistein or KBU 2046, or without treatment, compared to before treatment (set before treatment as 1).

Based on this data, we further investigated whether such methodology can be used to evaluate the effects of anti-metastatic drugs that target cellular motility, namely “migrastatic” compounds^2, 37^, on cancer cells. To this end, we tested the effects of two compounds known to inhibit the motility of prostate cancer cells^20, 21^, genistein and KBU 2046, on micromotion. Specifically, PC-3 cancer cells were encapsulated in fibrin gel and equilibrated in an incubator for 6h, then the culture medium in the 3D culture chamber was replaced with fresh medium (control), the medium containing either genistein or KBU 2046. Micromotion indexes were measured before, 24h, and 48h after treatment. The data for PC-3 cellular MIs treated by genistein or KBU 2046 are presented in Fig. 5G and H and control in Fig S4, respectively. Significant reductions of PC-3 cellular MIs were observed in the genistein and KBU 2046 treatment groups but was not observed in control, indicating that the reduction of MIs were induced by migrastatics rather than by other environmental factors. Interestingly, the micromotion variation dynamics of these two compounds are different. Significant reduction of MI was observed after 24h of genistein treatment (0.158 ± 0.053 vs 0.139 ± 0.058, P<0.05), but MI recovered back to the before treatment level (0.158 ± 0.053 vs 0.154 ± 0.077) after 48h. In contrast, KBU 2046 continuously reduced the MIs of PC-3 over the entire duration of the experiment. Although the inhibition effect of KBU 2046 to MIs at 24h after treatment is not significant (0.165 ± 0.071 vs 0.145 ± 0.045, P = 0.06), a reduction in MI was observed (0.165 ± 0.071 vs 0.133 ± 0.046, P < 0.01) after 48h KBU 2046 treatment with higher significance compared to genistein induced MI reduction. Figure 5I plots the fold changes of mean MIs of PC-3 at different time points with or without KBU-2046 or genistein treatment. The value in each square equals to the ratio of mean MIs between at certain time points and at before treatment ( e.g. 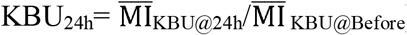). This plot illustrates the trend that KBU2046 and genistein exert similar effects at 24h but distinctive at 48h, in which time point the MIs of genistein treated group recovered to before treatment level while that of KBU treatment group reduced further. This set of data, particularly the recovery of MIs at 48h after genistein treatment, might also suggest PC-3 are of higher tolerance to genistein compared to KBU 2046. This is consistent with the observation from Li et. al that KBU 2046 is more effective in inhibition metastasis of prostate cancer cells than genistein^20^.

## Discussion

Tumour cell invasion of nearby tissues is one of the most important processes during cancer progression. Therefore, the identification of specific factors that can activate/suppress tumour cell invasiveness has strong potential to provide not only an improved mechanistic understanding of the metastatic process but also to assist in the development of novel therapeutic approaches. Importantly, solid tumours are inherently heterogeneous, being comprised of cellular subpopulations with vast differences in gene and protein expression, tumour-forming ability, and invasiveness^38–40^. In addition, a defining characteristic of tumour cells invasiveness is the fact that it inherently occurs in a 3D environment and over a relatively long period of time. Current approaches to determine the motility of tumour cells typically involve measuring their actual movement in space, which suffer from several limitations, especially when performed in a 3D environment^7–9^.

We report here a new paradigm, which is that measurement at the single cell level of an optical cellular micromotion index calculated from the dynamic nanoscale fluctuations of the OT is strongly correlated to intracellular biological processes underpinning cellular motility through ECM-like gel. The practical implementation of this concept enables the measurement, within a few minutes and non-invasively, of the invasiveness of single tumour cells within a 3D environment that mimics that of real tumour. The concept of cellular micromotion was initially demonstrated using impedance-based measurement of the nanoscale fluctuations of the distance between cell membranes and a gold electrode. Impedance based micromotion could be used to identify distinctive micromotion features of cancerous cells from normal cells. In addition, tumour cell lines with higher invasiveness were also shown to have higher micromotion (e.g. MDA-MB-231 vs MCF-7)^15^ This is in agreement with our observation that the optical cellular micromotion indexes of MDA-MB-231 cells were significantly higher than that calculated for the less invasive MCF-7 cells (0.170±0.06 vs 0.128±0.04, p< 0.01** at 6h after seeding and 0.165±0.05 vs 0.113±0.04, p< 0.001*** at 24h after seeding, Table. S1). However, impedance-based micromotion measurement is limited by the inherent requirement for the cells to be adhered on an aberrantly rigid electrode. In addition, while single cell measurements are theoretically possible, practical implementation requires the formation of a cellular monolayer onto the electrode and therefore the correlation of micromotion and invasiveness at the single cancer cell level has never been shown. While the impedance-based micromotion is an indirect measurement of the dynamic fluctuations of the cell membranes, the optical cellular micromotion as defined in this report is based on directly measuring the dynamic dry mass reorganisation that occurs within live cells. This reorganisation is measured as spatiotemporal fluctuations in the local OT (proportional to the cellular dry mass)^41^ which can be mapped with digital holographic microscopy in a high precision manner. Several previous studies reported that the cellular dry mass redistribution is driven by certain receptor activation^42, 43^. In particular, EGF stimulation was reported to mediate dynamic mass redistribution in epidermal growth factor receptor (EGFR) positive cancer cells^44, 45^. This is consistent with our observation that EGF upregulated the MI and migration capability of MDA-MD-231, although the cellular environment, as well as the objectives and analysis tools are very different between these two studies.

The “micromotion” as defined and measured in this study likely represents a composite of multiple intracellular processes that manifest as a single optically detectable cellular phenomenon, and presumably some cellular processes included in the observation and quantification are either less relevant or even irrelevant to cell migration/invasion. This “black box” aspect is the limitation of most label free observations^42^. To better interpret the measured “optical cellular micromotion”, the key regulators of micromotion and the mechanism behind such correlation between optical cellular micromotion and invasion were explored in this study. The key hypothesis here is that one or several intracellular machineries that drives cellular invasion overlaps with the optical micromotion measurements. It is well established that tumour cells actively migrate within tissues through dysregulation of cytoskeletal components such as actin dynamics and organization^46–48^. Our mechanistic data confirmed that the actin cytoskeleton is a significant regulator of optical cellular micromotion, supporting our speculation with respect to the mechanism behind the correlation.

Interestingly, we found that ~5 min of micromotion measurement can effectively distinguish cancer cells with different motility/invasiveness. This time span is much shorter than that of standard invasion assays based on measuring accumulative effects of invasion, such as distance or porous membrane translocation. This is not surprising, especially considering the dynamics of cytoskeleton activity in 3D environment. In fact, cytoskeletal dynamics that directly or indirectly link to cell invasion occur within similar time scale, including the displacement of actin filament (136s in a migrating keratocyte^49^) or microtubule redistribution (120s in human fibrosarcoma cells embedded in collagen I matrices^50^). A typical example is the dynamics of cellular protrusion that plays a crucial role in mediating ECM degradation and tumour cell invasion, such as lamellipodia, invadopodia or bleb. The formation and translocation of these actin and/or other cytoskeleton components abundant subcellular structures involve a series of transient, localized remodelling of the cell membrane and intracellular structures. For example, the formation of invadopodia involves vast cell membrane ruffling, extension and protrusion with only a few minutes lifetime^51^. These processes eventually result in rapid cell morphological changes^52–54^ and associated cellular dry mass redistribution.

It also worth noting that cells in the 3D environment were found to have significantly higher micromotion index than that in 2D (Fig. 2C). This is not surprising when considering the inherent differences between 2D and 3D environments, which change the modes of cell-matrix interaction and in turn influence cellular behaviours. For example, focal adhesions of cells growing in a 2D setting are substantially larger in scale (~15 μm vs ~ 0.3 μm) and more stably formed (~15 min vs ~1s of lifetime) than in 3D^6^. This distinction not only dictates the different motility of cancer cells^3^, but also could contribute to the substantial difference in micromotion for cells in 2D vs 3D observed in this study. Indeed, the substantially shorter lifetime of focal adhesion suggests that subcellular structures are reorganized far more rapidly in a 3D environment, which is expected to yield higher amount of micromotion within a period of time than cells in 2D.

One important aspect of the micromotion concept is its ability to inform about the heterogeneity within the tested population with respect to invasiveness as well as invasive phenotype (Fig. 3E and 4D). Tumours display high level of heterogeneity and potentially develop resistance to therapeutics with different mechanisms and in different degrees^31^. This heterogeneity can present in individual level, organ level (primary tumour vs metastasis), or intra-tumour level^55^. Therefore, indication of the killing (e.g. cytotoxic) or inhibition (e.g. cytostatic or migrastatic) effects towards such small portion of cells within the population is important in anti-tumour therapeutics screening and validation^40, 56^. Current population-based invasion measurements are often blind to important cellular subpopulations with high invasiveness, which likely negatively impact on their relevance to the *in vivo* situation, and the capabilities to evaluate the effectiveness of migrastatic compounds.

Being able to evaluate migrastatic drug efficacy is a key advantage of optical cellular micromotion. In contrast to conventional cytotoxic or cytostatic drugs that targeting the viability or growth of cancer cells, migrastatic drugs target the migration, invasion, and metastasis of cancer cells^2^. By specifically targeting cytoskeletal components of cancer cells, the small molecules disrupt machineries that play key roles in cancer cells invasion (e.g. cytoskeletal dynamics, lamellipodia/invadopodia formation/deformation, cell contraction/cell rear retraction)^57^. One example is KBU 2046, which decreases the motility of prostate cancer cells by modulating the activity of the oncoprotein HSP90. Our micromotion measurements support the hypothesis that KBU 2046 inhibits prostate cancer cell motility through disrupting the cytoskeleton reorganization, considering the association between micromotion and cytoskeleton activities. Also, optical cellular micromotion measurements provide insights into migrastatic compounds time response on both motility and cytoskeleton activity of single cells. Taken together our results show the feasibility of using optical cellular micromotion measurements to assess the efficacy of candidate migrastatic drugs. It also suggests that it could be used to monitor transient invasiveness changes of cancer cells at specific time point during drug treatment, which is not possible in standard invasion assay but could be very useful towards yielding better understanding of the mechanisms underpinning cancer motility inhibition.

A number of limitations in this study should be considered and warrant future investigation. First, although cells were suspended in fibrin gel that support cellular motility in 3D, the ECM biological and structural complexity (e.g. complex ECM bundles, spatial organization of ECM fibers) is not fully recapitulated by fibrin. The specific structure of the ECM regulates tumour cells motilities through driving the dynamic re-organization of cytoskeleton and focal adhesion.^58, 59^ Micromotion indexes measured in more complex gels that better mimic the native ECM of specific tissues is therefore anticipated to be of higher physiological relevance and consequently could yield better biological insight. In addition, only isolated single cells with clear cell boundaries were included in the micromotion measurements in the present study as this significantly reduces the technical difficulties in image acquisition and processing (e.g. image segmentation). However, cell-cell interactions play a significant role in the migration of cells through tissues^60, 61^ and refinement of the method to enable such complex measurement could further strengthen its significance.

## Materials and methods

### Materials

Phosphate buffer saline (PBS), 4% formalin solution, fibrinogen and thrombin from bovine plasma were all purchased from Sigma-Aldridge. Poly(dimethylsiloxane) (PDMS) was acquired from Dow Corning (Singapore). Cytochalasin D was also obtained from Sigma-Aldridge and dissolved in DMSO as stock solution. DMEM (pH = 7.4, Sigma-Aldridge) for MDA-MB-231 and MCF-7 breast cancer cell lines culture and F-12K (ATCC) for A549 VIM-RFP culture were supplemented with 10 % fetal bovine serum (FBS, Gibco) solution and 1 % antibiotics (Gibco). Both normal PC-3 prostate cancer cell line (ATCC) and PC-3 transfected with miR-194 or negative control mimic were maintained in RPMI-1640 medium (Gibco) supplemented with 10 % fetal bovine serum (FBS, Gibco) solution and 1 % antibiotics (Gibco). Human umbilical vein endothelial cells (HUVECs) and normal human lung fibroblasts (NHLFs) were purchased from Lonza and cultured in EGM-2 and FGM-2 (Lonza) supplemented with EGM-MV and FGM-2 Bullet Kit, respectively. Human epidermal growth factor (hEGF) was purchased from Lonza and EMT inducing media supplement was purchased from R&D System.

### Cell Culture

MDA-MB-231, MCF-7 breast cancer cell lines, A549 VIM-RFP cancer cell line and PC-3 prostate cancer cell line with or without transfection (ATCC) used in this study were cultured in 75mm^2^ tissue culture flasks and maintained in 37 °C and 5% CO_2_ humidifying incubator. Medium was changed every 2-3 days and cells were subcultured until reach 70-80% confluency. For DHM imaging, a custom-made device (imaging device) was used to culture cell both in 2D and 3D. A PDMS chamber with 2 openings for injecting medium was made with both sides sealed by optically clear polymer coverslips (Ibidi), in order to make both sides flat and optically transparent for DHM imaging. The distance between the two coverslips was 300 μm. For 2D cell culture, the cells were seeded directly into the chamber after both sides have been sealed. A tissue culture treated polymer coverslip at the bottom was used. For 3D cell culture, fibrin gel was used for extracellular matrix. Only one side of chamber was sealed with a coverslip and the other side was left unsealed before cell seeding. Subsequently 5 μL of cell suspension with 5U/mL thrombin and 5 μL of 10 mg/mL fibrinogen solution was homogeneously mixed. 5 μL of the cell/fibrin gel mixture was dropped onto the coverslip in the PDMS chamber which was sealed with another coverslip. After 10 minutes when the fibrin gel completely gelled, cell culture medium was injected into the PDMS chamber. In both experiments the final concentrations of cells (in medium, or in pre-gel solution) were 5 × 10^6^/mL.

### Micromotion index measurement

In this study, a commercially available DHM® T-1000 (Lyncée Tech) was used. The custom-made device for cell culture was kept in a stage-top incubator (Chamlide) maintained at 37 °C and 5% CO_2_ during the real time DHM imaging. Koala software native to the DHM was used to control the imaging and reconstructed holographic images to phase images. The software allows automated conversion of raw phase images into OT measurements. It is worth noting here that practically what is being measured with the DHM is the phase shift of the light that passes through a cell but it is proportional to the cell optical thickness (OT). In this sense we refer to “phase shift” when we discuss light or optics but “OT” when we refer to cells.

For micromotion measurement, 1024 images were typically acquired every 250 ms with the 20× magnification objective. In the case of 3D culture, each cells were digitally focussed by adjusting the “focus” parameter from “reconstruction settings” window in the Koala software. In this case new region of interest (ROI) was defined and the in-focus cell was included in the reconstructed phase images. Reconstructed phase images with single focused cell rather than raw phase images that include both in-focus and out-of-focus cells, were used for micromotion measurement in the next step and were presented in the figures. Since we were interested in the optical cellular micromotion of the single cells, in the study only individual cells with clear cell boundary were included in the analysis. Clusters of cells or cells overlapped with others were removed from analysis. In addition, in 3D cell measurement, cells with average OT lower than average value of 2D cells and has typical 2D morphology were considered as 2D cells and were removed from analysis.

A mask for each cell is first built to segment cells from the background automatically by an algorithm using a threshold determined from pixel height. We used the method reported by Shaked et. al to quantify the dynamic OT (OT) fluctuation profile. In brief, we define the shift of OT (ΔOT) using eq (1):

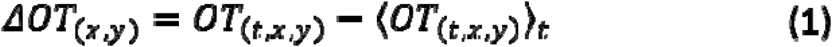

in which OT_(t,x,y)_ refers to spatially variant OT of pixel (x,y) at time point t and 〈OT_(t,x,y)_ 〉_t_ refers to the average of OT_(t,x,y)_ over the entire imaging period. Then we calculate the mean square of ΔOT using eq (2):

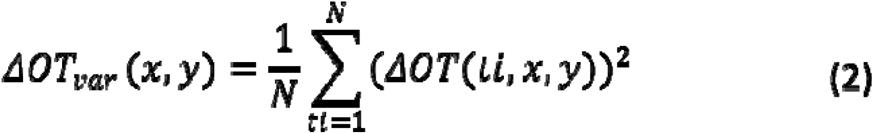

in which i=[1,…, 1024] is indexing the number of images. Lastly we define the MI with eq (3):

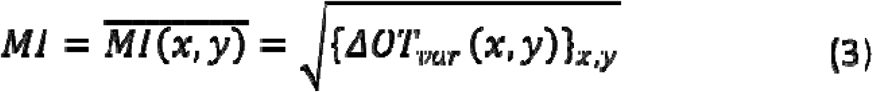

in which the MI of every pixel in a cell mask is calculated then averaged over the whole mask

### Cell random migration assay and speed calculation

MDA-MB-231 breast cancer cells were used for random migration assay. After 5 min imaging with DHM to obtain the MIs, a 12 h DHM imaging was carried out, in which 72 images were acquired every 10 min using the 10× magnification objective. The Trackmate plugin in open access software Fiji was then used to automatically segment and track the cell movement and record the position of cell centre (x,y) in every frame. The mean displacement between each frame was calculated with eq (4)

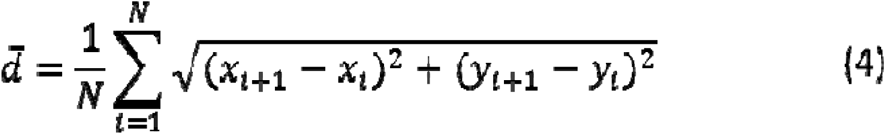

in which i=[1,…,72] is indexing the number of images. Mean speed (v) equals to the /frame.

### Actin cytoskeleton inhibition

Both MDA-MB-231 and PC-3 were used for action cytoskeleton inhibition assay. Actin cytoskeleton inhibition was performed directly in 3D environment. In specific, 1 mg/mL cytochalasin D stock solution that dissolved in DMSO was 1:200 diluted in 6 mg/mL fibrinogen solution, which then quickly mixed with same volume of cell suspension and seeded in the device. The cell seeding procedures followed the 3D cell culture protocol described in previous cell culture section. In this case the 3D fibrin gel was supplemented with 2.5 μg/mL cytochalasin D. In the control group DMSO at final concentration of 0.4% (v/v) was supplemented. Then device was filled up with cell culture medium containing 2.5 μg/mL cytochalasin D or 0.4% DMSO and kept in incubator for 2h for equilibration before micromotion measurement.

### EMT induction

MCF-7 and A-549 was used for EMT induction assay. EMT induction was performed on 2D according to the protocol provided by manufacturer, then encapsulated in gel for 3D measurements. Briefly, 1×10^5^ cells were seeded in 6 well plate with 2 mL of the culture medium supplemented with 1% StemXVivo EMT inducing media supplement. The medium was replaced with fresh EMT induction medium on day 3 after induction. After 5 days the EMT induction is completed and cells are used as described above for DHM imaging. Imaging of VIM-RFP reporter was performed using LSM 710 confocal microscope (Zeiss, Germany) and quantification of fluorescent intensity of single A549 VIM-RFP cell was performed using ImageJ (NIH). Single fluorescent cells were segmented from background manually for determining of area (μm^2^) and fluorescence intensity (F.I.) was measured automatically by ImageJ. Mean fluorescence intensity (M.F.I.) was calculated using F.I divided by area of single cells (M.F.I.= F.I. / area).

### MiRNA transfection and intravenous metastasis assay

Both the transfection and intravenous metastasis assay were conducted following the protocols reported previously^19^ without any modification. Intravenous metastasis assay were approved by the University of Adelaide Animal Ethics Committee (approval number M-2014-180C).

### Microvasculature-on-a-chip preparation

The preparation of microvasculature-on-a-chip was performed following the protocol developed by Chen et. al^35^ with slight modification as we previously reported^62^. Briefly, HUVECs (2.5×10^7^/mL) and NHLFs (1.2×10^7^/mL) were suspended in endothelial culture medium supplemented with 4 U/mL thrombin and mixed with equal volume of 6 mg/mL fibrinogen solution. After quickly pipetting, cells-gel mixture was injected into the corresponding cell channel and incubated by approximately 10 min for the gelation of fibrinogen. Medium in the device was changed every day until the formation of well-formed microvasculature (approximately 5 days).

### Extravasation assay

The tumour cell extravasation assay was performed after 5 days when the microvasculature is well-formed and more than 50% of interpost regions (regions in between the device micropillars) have established vascular openings. For extravasation assay, the medium was withdrawn from all the reservoirs and a 50 μL of control or miR-overexpressed PC-3 cells suspension with a concentration 2×10^5^ /mL was pipetted into one reservoir linked to the medium channel that flanks the HUVEC (microvasculature) channel. The medium was allowed to perfuse across the medium channel gradually to fill up the reservoir on the other side of medium channel, driven by hydrostatic pressure. After reaching equilibrium of hydrostatic pressure, another 50 μL of cell suspension was added into both reservoirs on one side to establish a hydrostatic pressure across the microvasculature channel and the device was placed in the incubator. After 10 min, when cancer cells were fully perfused across the microvasculature, the cell suspension was withdrawn, and fresh medium was added to all the reservoirs to wash away cancer cells not attached to the medium channel. Finally, the number of perfused and arrested cells were observed under bright field microscopy to ensure there are enough cells perfused and arrested into microvasculature (typically around 100-200 cancer cells per device). The extravasation rate was calculated as the proportion of cancer cells that escaped from the microvasculature relative to the total number of cells that remained inside the microvasculature on chip device. Identification of cancer cells that escaped from microvasculature was performed using confocal microscopic images (LSM 710 confocal microscope). Boundary of microvasculature was visualized by endothelial marker (VE-cadherin) labelled with standard immunostaining method, PC-3 with or without overexpression of miR-194 were visualized by self-tagged fluorescence.

### “Migrastatic” compound treatment

KBU 2046 is kindly provided by Raymond Bergan’s laboratory and preparation of KBU 2046 cell treatment solution follows the protocol provided by Raymond Bergan’s laboratory. Briefly, 5 mg of KBU 2046 was dissolved in 206.4 μL to prepare 100 mM stock solution, then aliquoted 10 μL such solution in 1.5 mL Eppendorf tube and preserved in −20□. Before use, stock solution was kept in 37□ incubator until the solution was completely thawed and 90 μL additional DMSO was added into tube to prepare 10 mM solution. Then the solution was 1:1000 diluted in cell culture medium to obtain 10 μM working solution. Both 10 mM DMSO solution and 10 μM working solution will be disposed immediately after experiment and will not be reused. Genistein was purchased from Sigma and similar protocol was followed to prepare both stock and working solution with only modification that dissolving 5 mg genistein in 185.0 μL DMSO to prepare 100 mM stock solution. For in-device cell treatment experiment, cells were seeded in fibrin gel in device and incubated for 6h before DHM imaging. After imaging completed, the device was placed back to incubator for at least 15 min before performing medium replacement. For the medium replacement, a drop of fresh medium (with or without drugs) was placed on top of both inlet and outlet with 1 mL pipette to wet the PDMS surrounding the inlet and outlet, followed by removing most of medium on outlet and adding medium on inlet. Driven by hydrostatic force, the flow of medium from inlet to outlet will form, and then keep removing medium on outlet to continue the flow until most of medium has flowed through the device. Repeat such procedures at least twice to make sure most medium in the chamber has been replaced with fresh medium.

### Statistical analysis

Statistical analyses were performed using student t-test for any data set comparison. P< 0.05 was considered to be statistically significant.

## Acknowledge

This work was performed in part at the South Australian node of the Australian National Fabrication Facility, a company established under the National Collaborative Research Infrastructure Strategy to provide nano and microfabrication facilities for Australia’s researchers. We thank Prof. Raymond Bergan and his laboratory from Oregon Health & Science University to provide KBU 2046 compound and relevant documents. This work was supported by the ARC Centre of Excellence in Convergent Bio and Nano Science and Technology and funding from the National Health and Medical Research Council of Australia (ID 1083961 to LAS). LAS is supported by a Principal Cancer Research Fellowship awarded by Cancer Council’s Beat Cancer project on behalf of its donors, the state Government through the Department of Health and the Australian Government through the Medical Research Future Fund.

