## Supplemental information for "Optical Cellular Micromotion: A New Paradigm to Measure Tumour Cells Invasion in 3D Tumour Environments"

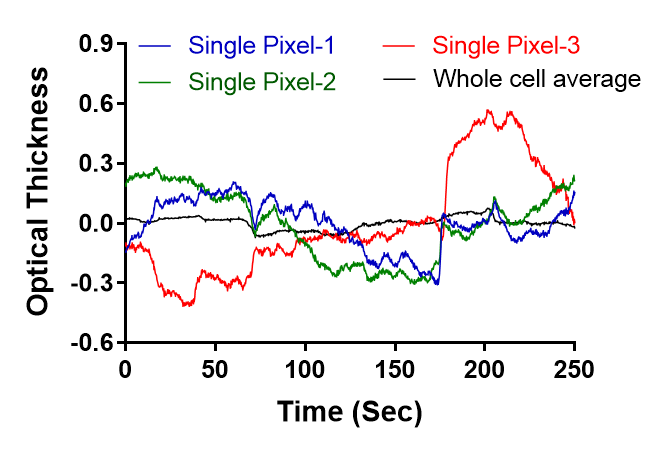

**Supplementary Figure 1.** Fluctuation curves of the OT averaged for a whole cell (black) and of the OT for 3 random single pixels (blue, red and green) inside the same cell.

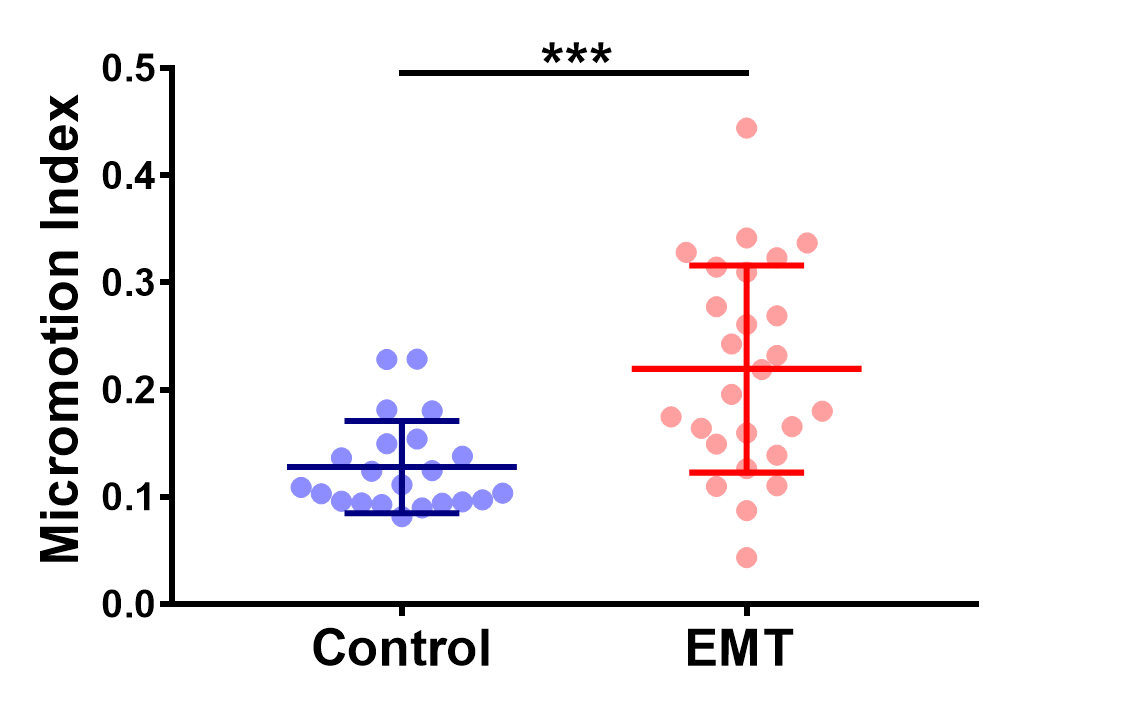

Supplementary Figure 2. MI for MCF-7 with and without EMT induction. Error bars are SDs (***, P< 0.001 unpaired two-sided t-test).

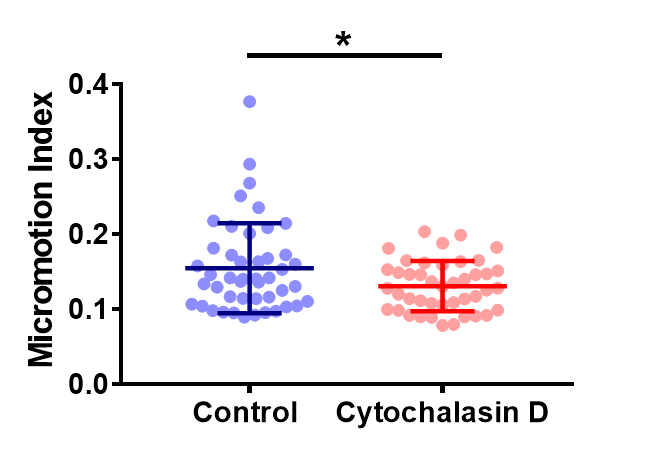

Supplementary Figure 3. MI for PC-3 with and without Cytochalasin D inhibition. Error bars are SDs (*, P< 0.05 unpaired two-sided t-test).

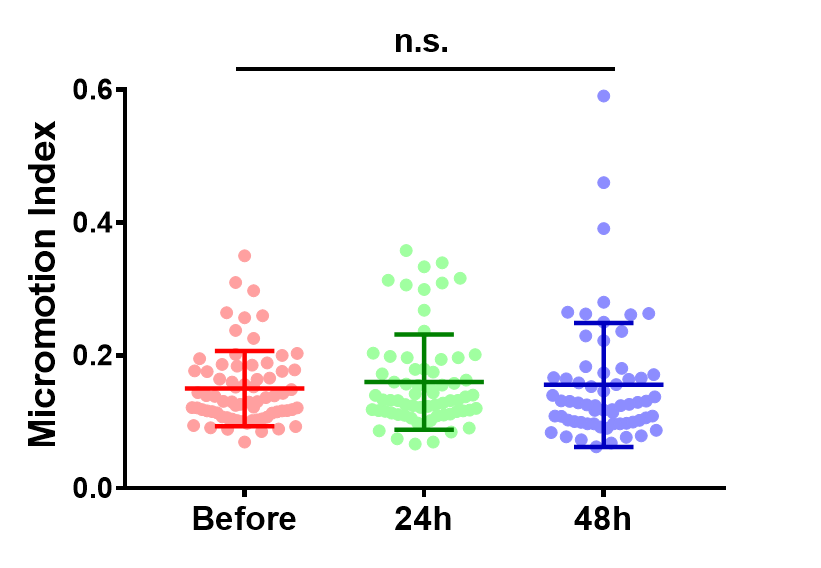

Supplementary Figure 4. MI for PC-3 in the micromotion measurement device without migrastatic compound treatment (control group). Before, 24h, and 48h indicate the measurement time points that consistent to the treatment groups. Error bars are SDs (*, P< 0.05 unpaired two-sided t-test).

|  | 6h after seeding | 24h after seeding | P value (6h vs 24h) |
| --- | --- | --- | --- |
| MDA-MB-231 | 0.170 ± 0.06 | 0.165 ± 0.05 | 0.696 (n.s.) |
| MCF-7 | 0.128 ± 0.04 | 0.113 ± 0.04 | 0.261 (n.s.) |
| P Value (MDA-MB-231 vs MCF-7) | 0.0051** | 0.0001*** |  |

Supplementary Table 1. MIs calculated for MDA-MB-231 and MCF-7 cells at 6h or 24h after being seeded into 3D fibrin gels (unpaired two-sided t-test).

**Supplementary Movie 1** Representative cellular OT fluctuation movie in 3D environment. (heat map represents cellular optical thickness and was scaled automatically)

**Supplementary Movie 2** Representative cellular OT fluctuation movie in 2D environment. (heat map represents cellular optical thickness and was scaled automatically)

**Supplementary Movie 3** Cellular OT fluctuation movie of the representative single MDA-MB-231 in 3D environment with exogenous EGF. (Movie plays at 25 × real speed)

**Supplementary Movie 4** The migration movie of the cell demonstrated in Movie S3 in following 12h after micromotion imaging.

**Supplementary Movie 5** Cellular OT fluctuation movie of representative single MDA-MB-231 in 3D environment without exogenous EGF.

**Supplementary Movie 6** The migration movie of the cell demonstrated in Movie S5 in following 12h after micromotion imaging.
